# Senescent cells are more susceptible to reductive stress-induced cell death: implications for senolytic research

**DOI:** 10.64898/2026.07.10.737740

**Authors:** Vladimir Belhac, Alexandra Stolzing, Neil R.W. Martin

## Abstract

Proliferating cells can enter an irreversible state of cell-cycle arrest known as cellular senescence. The accumulation of senescent cells contributes to organismal ageing and age-related pathologies. Consequently, therapeutic strategies have emerged to selectively eliminate senescent cells (senolytics). Our previous work suggested that senescent mouse myoblasts are more susceptible to reductive stress-induced cell death than proliferating cells. Here, we replicated these findings in human LHCN-M2 myoblasts, demonstrating a biphasic dose–response relationship with cell death, wherein low concentrations were associated with reduced cell death in both proliferating and senescent cells, whereas higher concentrations selectively induced cytotoxicity in senescent cells. We propose that many identified natural senolytic compounds may exert their in vitro activity, at least in part, through the induction of reductive stress due to their antioxidant properties. These findings have important implications for understanding senolytic mechanisms and guiding the future development of senescence-targeting therapies.

## Main text

A notable aspect of mammalian ageing has been the accumulation of senescent cells. Mitotically competent cells adopt a senescent state in response to replicative exhaustion, sublethal stress, or activation of oncogenes. This senescent state is characterised by a set of commonly observed hallmarks, including cell cycle arrest, a pro-inflammatory secretome, organelle dysfunction, and, notably, resistance to cell death (Hernandez-Segura, Nehme and Demaria, 2018; Di Micco *et al*., 2021).

Senescent cells play a dual role in biology. On one hand, they contribute to normal physiological processes such as wound healing and development (de Magalhães, 2024). On the other, their accumulation is increasingly recognised as a contributor to organismal ageing. Senescent cell burden increases markedly during ageing (Herbig *et al*., 2006), with human tissues from older donors (> 66 years) showing up to tenfold higher levels of p21- and p16-positive cells, markers of senescence-associated cell cycle arrest, in comparison to young donors (≤ 35 years) (Idda *et al*., 2020). Moreover, genetic elimination of senescent cells in mice, achieved through inducible apoptosis of p16-expressing cells, led to extended lifespan and improvements in multiple ageing phenotypes, demonstrating a causal role for senescence in the ageing process (Baker *et al*., 2011). Subsequently, there has been a drive to identify compounds capable of selectively inducing death in senescent but not proliferating cells, termed senolytics. Among the most well-characterised senolytics are BCL-2 family inhibitors, such as Navitoclax, and tyrosine kinase inhibitors, including Dasatinib (Zhu *et al*., 2015; Kirkland and Tchkonia, 2020; Di Micco *et al*., 2021).

Another class of senolytics includes natural compounds, particularly flavonoids such as Fisetin, which is currently undergoing a clinical trial for its senolytic properties (Ji *et al*., 2026). Although precise mechanisms remain undefined, the effects of such compounds are often attributed to inhibition of the PI3K/Akt/mTORC1 signalling pathway, which senescent cells are more dependent on for survival (Di Micco *et al*., 2021; Tavenier *et al*., 2024). Our recent work demonstrated that an acute bolus of synthetic antioxidant suppresses PI3K/Akt/mTORC1 signalling (Belhac *et al*., 2026). Such a response is not unheard of, since ROS has been implicated in the regulation of upstream kinases and phosphatases through reversible oxidation of cysteine thiols (Sies and Jones, 2020). Interestingly, flavonoid-based senolytics also possess potent antioxidant properties (Rolt and Cox, 2020), raising the possibility that their inhibitory effects on PI3K/Akt/mTORC1, and subsequent senolytic activity, are mediated through redox modulation. We confirmed these observations in hypothesis-driven experiments demonstrating that, following prolonged exposure to synthetic antioxidants, senescent C2C12 mice myoblasts were more prone to antioxidant-induced cell death compared to proliferating myoblasts (Belhac *et al*., 2026).

Here, we investigated whether these findings could be replicated in human myoblasts and further characterised the antioxidant’s cytotoxic effects using dose–response experiments. Senescence was induced in LHCN-M2 myoblasts by treatment with 200 nM doxorubicin for 2 days, followed by an 8-day recovery period before a 3-day exposure to the indicated treatments. We show that (i) a cytotoxic concentration of DMSO induced a nearly twofold greater increase in relative cell death in proliferating than in senescent cells (Figure 1A), consistent with the well-established resistance of senescent cells to cell death (Hernandez-Segura, Nehme and Demaria, 2018; Di Micco *et al*., 2021); and (ii) the antioxidant N-acetylcysteine (NAC) displayed a biphasic dose–response relationship with cell death, with low concentrations tending to reduce cell death in both proliferating and senescent cells, whereas higher concentrations selectively induced cytotoxicity in senescent cells (Figures 1B and C). Together, these findings suggest that, compared with proliferating myoblasts, senescent human myoblasts are more susceptible to reductive stress-induced cytotoxicity.

**Figure 1.**
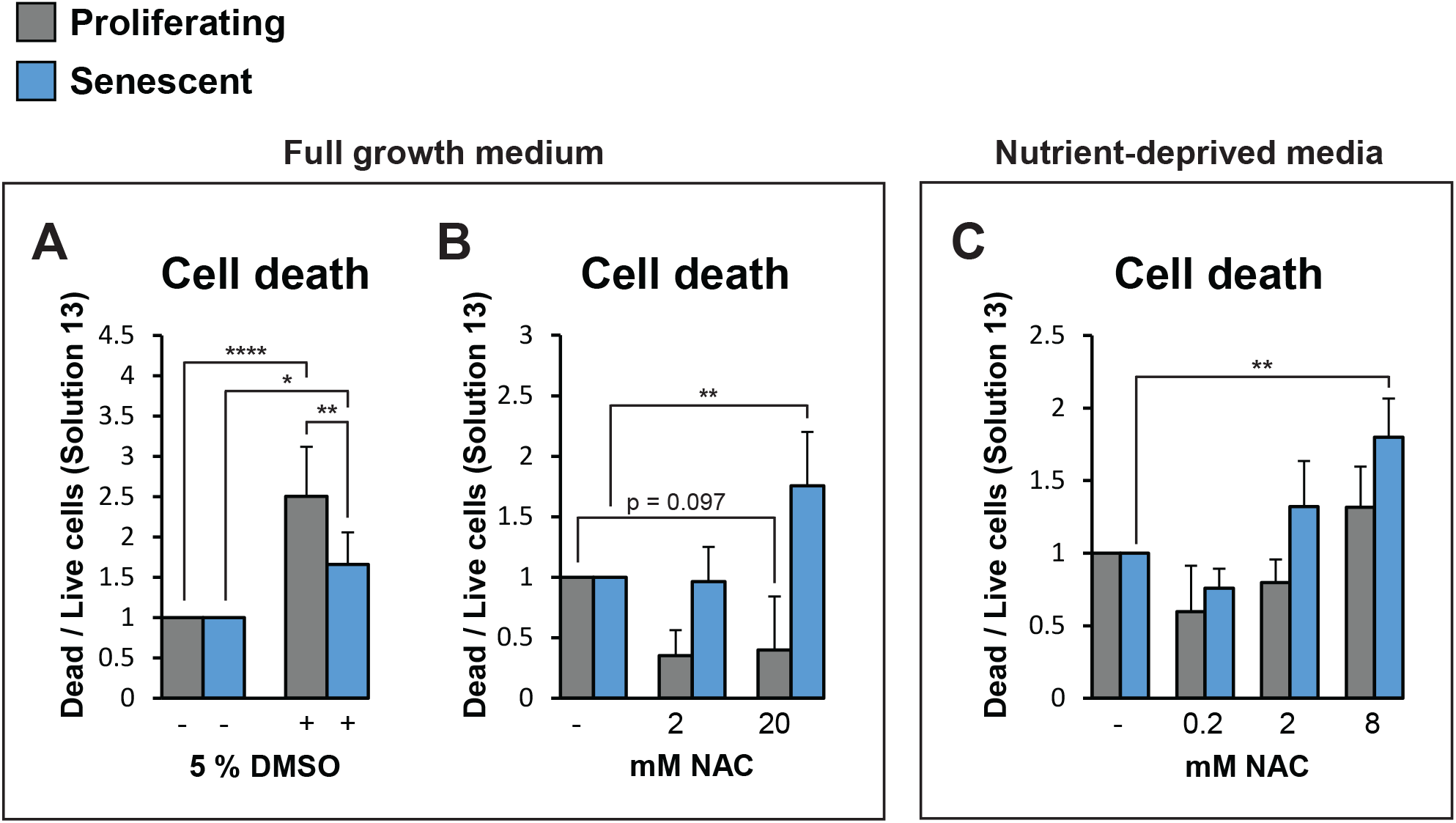
Senescent myoblasts are more susceptible to reductive stress-induced cell death. Doxorubicin-induced senescent and proliferating myoblasts were treated as indicated for three days in either full growth medium or amino acid-, glucose-, and serum-restricted medium. (**A**) Exposure to a cytotoxic concentration of DMSO resulted in a higher proportion of dead cells in proliferating than in senescent cultures, consistent with the previously reported resistance of senescent cells to cell death. (**B** and **C**) Exposure to increasing concentrations of NAC produced the opposite effect to that observed in (**A**), with higher NAC concentrations increasing the proportion of dead senescent cells. Results are expressed as relative values normalized to untreated cells and are presented as mean ± standard deviation. P values were calculated using two-way ANOVA followed by estimation of marginal means with Bonferroni correction. *P ≤ 0.05, **P ≤ 0.01, ****P ≤ 0.0001. n = 3. DMSO,

Although suppression of PI3K/Akt/mTORC1 signalling remains a plausible explanation for the senolytic properties of antioxidants described above, we suspect that reductive stress may also contribute, either as an alternative or complementary mechanism. Often described as the counterpart to oxidative stress, reductive stress is characterised by an excess of reducing agents such as reduced glutathione and NADPH (Pérez-Torres, Guarner-Lans and Rubio-Ruiz, 2017; Kunwar and Aishwarya, 2024). There is some evidence to suggest that senescent cells may be more susceptible to reductive stress. While they are commonly characterised by elevated ROS production, studies have reported lower ROS levels within the endoplasmic reticulum (ER) compared to proliferating cells (Qiao *et al*., 2022). This is notable because, compared with the cytosol, the ER relies on orders of magnitude higher ROS levels to facilitate protein folding (Sies and Jones, 2020), and exacerbated ER stress and the resulting cell death has been described as key consequences of reductive stress (Pérez-Torres, Guarner-Lans and Rubio-Ruiz, 2017; Kunwar and Aishwarya, 2024). Taken together, these observations suggest that senescent cells may already exhibit some localised form of reductive stress at baseline, which could arguably make them more susceptible to further perturbation by the addition of exogenous antioxidant compounds. In support of this, our data showed that treatment of senescent cells with the antioxidant NAC further increased the already elevated expression of ER stress-related mRNAs (DDIT3 and XBP1) (Belhac *et al*., 2026). It should also be noted that reductive stress can be cytotoxic through other mechanisms. This includes their capacity to have localised, pro-oxidant counter responses (Pérez-Torres, Guarner-Lans and Rubio-Ruiz, 2017; Kunwar and Aishwarya, 2024). For example, enhancing glutathione reductive capacity through pharmacological (NAC) and genetic means was reported to increase mitochondrial ROS production and reduce cell viability (Zhang *et al*., 2012; Kolossov *et al*., 2015; Noch *et al*., 2024). Antioxidants are also reported to exhibit pro-oxidant effect in the presence of transition metals, such as iron (Halliwell, 1996; Sagristá *et al*., 2002), which may be particularly relevant given that senescent cells accumulate iron at substantially higher concentrations (Masaldan *et al*., 2018; Admasu *et al*., 2023). Altogether these findings demonstrate how reductive stress through synthetic or natural compounds can be cytotoxic and can be particularly prominent in senescent cells.

The biological link between the senolytic properties of flavonoids and reductive stress has not been previously acknowledged. One possible reason for this oversight is that the conclusion is counterintuitive, given that low/moderate doses of antioxidants generally increase cell viability, even in senescent cells (Khor *et al*., 2017) (also shown in Figure 1C, albeit not reaching statistical significance). Note, however, that this response is not specific to synthetic antioxidants but also to flavonoids (Agullo *et al*., 1996; Dong *et al*., 2025). We speculate that it is the high dose and/or prolonged treatment duration that facilitates the transition from cytoprotective to cytotoxic response through reductive stress. Based on our interpretations of previously published data, we believe that such a pattern is in fact observed in some of the seminal papers which demonstrated that the flavonoid fisetin is senolytic. In these studies, treatment of senescent cells with fisetin results in a biphasic relationship, whereby low fisetin doses appear to cause a mild increase in cell survival/number, while the higher concentration appears to decrease it (Zhu *et al*., 2017; Yousefzadeh *et al*., 2018). We further speculate that differences in antioxidant potency between flavonoids and synthetic antioxidants may explain why reductive stress is difficult to identify as a senolytic mechanism. For example, The Trolox Equivalent Antioxidant Capacity (TEAC) for fisetin and quercetin in vitro is ≥ 2.8 (Ishige, Schubert and Sagara, 2001), whereas the TEAC for NAC is ∼0.5 (Güngör *et al*., 2011). Consistent with this observation, NAC is typically used at concentrations of up to ∼5 mM, which are orders of magnitude higher than those used for flavonoids (Wu *et al*., 2014; Park *et al*., 2023). Consequently, to induce reductive stress with NAC would require concentrations greater than 10 mM which would then increase the risk of osmotic stress, and as such, these concentrations are generally avoided by scientists working in vitro. Our initial experiments have bypassed this fact by conducting these experiments in partially nutrient deprived condition, which in our experience, increase the susceptibility to cell death and decrease the concentration at which antioxidants become cytotoxic (see comparison between Figure 1B and C).

The senescent phenotype is highly heterogeneous, which may influence its susceptibility to reductive stress-induced cell death. One factor predicting this susceptibility could be a cell’s endogenous antioxidant activity or protein expression, which have been reported to increase (Khor et al. 2017; Borlon et al. 2007) or decrease (Yu *et al*., 2018; Lee *et al*., 2020) in senescent cells, depending on cell type and other experimental conditions. Assuming that our observations about reductive stress are true, this could explain some of the contrasting findings with flavonoids. For instance, one study reported that fisetin is senolytic in HUVECs (human umbilical vein endothelial cells) but not in primary human preadipocytes (Zhu *et al*., 2017). Therefore, the senescent cell heterogeneity may make reductive stress a senolytic property that is not universal.

Both flavonoids (Zhu *et al*., 2015; Yousefzadeh *et al*., 2018) and antioxidants (Breau *et al*., 2019; Wu *et al*., 2020; Sohouli *et al*., 2023) have been reported to reduce markers of senescence in vivo. However, it remains unclear whether these effects arise from attenuation of the senescent phenotype (senomorphic action) or their elimination (senolysis) since both antioxidants and senolytics more broadly, often have multiple biological targets (Kirkland and Tchkonia, 2015; de Magalhães, 2025). If reductive stress underlies senolysis, inducing sufficient reductive stress in senescent cells in vivo without detrimental systemic effects is unlikely without highly selective delivery approaches, such as those targeting senescent cell surface markers (Poblocka *et al*., 2021) or lysosomal β-galactosidase activity (Muñoz-Espín *et al*., 2018). On the other hand, our work shows that when the antioxidant NAC is used at low concentrations in vitro, it alleviates certain aspects of the senescent phenotype (Belhac *et al*., 2026). Moreover, antioxidants have been reported to prevent the acquisition of multiple features of senescence, including standard markers of cell cycle arrest (Zhang *et al*., 2015; García-Prat *et al*., 2016; Nacarelli, Azar and Sell, 2016; Yang *et al*., 2018; Summer *et al*., 2019). Therefore, it may be plausible that, in vivo, these compounds primarily act as senomorphics.

While we provide a justification for why the senolytic properties of some of the natural compounds are related to reductive stress, we do not exclude other possible mechanism that may act in unison. There is some evidence that flavonoids can bind to protein kinases such as receptor tyrosine kinases (Hou and Kumamoto, 2010), however the functional consequence remains unclear.

Overall, our work reveals that reductive stress, in addition to the more widely recognized oxidative stress, can drive cytotoxicity. Thus, it is the loss of redox balance, rather than oxidative or reductive stress alone, that appears to be critical. Pharmacological approaches that disrupt redox homeostasis in either direction may therefore exploit vulnerabilities in senescent cells to reduce their detrimental impact on tissue, organ, and organismal health.

## Methods

LHCN-M2 myoblasts (Evercyte) were cultured according to the manufacturer’s instructions. Cell culture dishes were pre-coated with 0.1% gelatin (Sigma-Aldrich), and the growth medium consisted of high-glucose Dulbecco’s Modified Eagle Medium (DMEM) supplemented with GlutaMAX, 20% Medium 199, 15% fetal bovine serum (FBS), and 1% penicillin–streptomycin (all from Gibco), 20 mM HEPES, 3 µg/mL zinc sulfate, 1.4 µg/mL vitamin B12, 0.055 µg/mL dexamethasone (all from Sigma-Aldrich), 2.5 ng/mL human growth hormone, and 5 ng/mL basic fibroblast growth factor (bFGF) (both from PeproTech).

For nutrient deprivation, cells were cultured in 30% growth medium diluted with amino acid-, glucose-, and serum-free DMEM (USBiological Life Sciences). Cellular senescence was induced by treating cells at ∼80% confluence with 200 nM doxorubicin (dissolved in DMSO; final DMSO concentration, 0.002%) for 2 days, followed by an additional 8 days of culture in doxorubicin-free medium.

Proliferating and senescent cells in 6-well plates were treated at 70% confluence with DMSO (Thermo Fisher Scientific) or N-acetylcysteine (NAC; Sigma-Aldrich) for three days in either growth medium or nutrient-deprived medium. Cells were then trypsinised, stained with Solution 13 (ChemoMetec) containing acridine orange (AO, which stains all nucleated cells) and DAPI (which stains cells with compromised plasma membranes), allowing discrimination between live (AO^+^/DAPI^−^) and dead (AO^+^/DAPI^+^) cells, and imaged using a NucleoCounter NC-3000 (ChemoMetec). Images were quantified manually in ImageJ using particle analysis. The macro used for analysis is available at: https://github.com/VladimirBelhac/ImageJ-macros. P values were calculated using two-way ANOVA followed by estimation of marginal means with Bonferroni corrections.

## Data availability

The data that support the findings of this study are available from the corresponding author upon reasonable request

## Acknowledgements

The authors sincerely thank Professor Lynn Cox (University of Oxford) for her valuable feedback on the manuscript.

